# Structure of the eukaryotic cytoplasmic pre-40S ribosomal subunit

**DOI:** 10.1101/216713

**Authors:** Alain Scaiola, Cohue Peña, Melanie Weisser, Daniel Böhringer, Marc Leibundgut, Purnima Klingauf-Nerurkar, Stefan Gerhardy, Vikram Govind Panse, Nenad Ban

## Abstract

Final maturation of eukaryotic ribosomes occurs in the cytoplasm and requires the sequential removal of associated assembly factors and processing of the immature 20S pre-RNA. Using cryo-electron microscopy (cryo-EM), we have determined the structure of a cytoplasmic pre-40S particle poised to initiate final maturation at a resolution of 3.4 Å. The structure reveals the extent of conformational rearrangements of the 3’ major and 3’ minor domains of the ribosomal RNA that take place during maturation, as well as the roles of the assembly factors Enp1, Ltv1, Rio2, Tsr1, and Pno1 in the process. Altogether, we provide a structural framework for the coordination of the final maturation events that drive a pre-40S particle towards the mature form capable of engaging in translation.

## Introduction

The ribosome performs the essential task of decoding genetic information into proteins. While structures of the mature ribosomal subunits in different functional states provide detailed insights into the mechanisms of translation, our knowledge regarding ribosome assembly and quality control at atomic resolution is only emerging.

In eukaryotes, ribosome assembly is a complex process that stretches across the nucleolus, nucleoplasm and cytoplasm and requires the concerted action of >200 assembly factors and several energy-consuming enzymes (Konikkat & Woolford, 2017, Kressler, Hurt et al., 2017, Pena, Hurt et al., 2017, Woolford & Baserga, 2013). The process is initiated in the nucleolus with the RNA Pol-I driven synthesis of the 35S pre-rRNA. The emerging pre-rRNA undergoes co-transcriptional folding and modification and recruits ribosomal proteins (r-proteins) and assembly factors specific for the small ribosomal subunit to form the earliest ribosome precursor, the 90S. Cleavage within the pre-rRNA releases the earliest small ribosomal subunit 40S precursor (pre-40S). Pre-60S large ribosomal subunit and pre-40S particles mature independently by transiently interacting with assembly factors (AFs) and are separately transported by export receptors through the nuclear pore complexes into the cytoplasm. Cytoplasmic maturation events are intimately intertwined with quality control and proofreading of functional centers, which ensures that only correctly assembled ribosomal subunits are committed to the translating pool (Greber, 2016, Greber, Gerhardy et al., 2016).

Exported pre-40S ribosomal particles undergo a series of cytoplasmic maturation steps before engaging in translation, including the sequential release of assembly factors and export receptors, as well as the assembly and stable incorporation of remaining ribosomal proteins. During the final steps of cytoplasmic maturation of the pre-40S, the 3’ end of the 20S rRNA is cleaved off by the nuclease Nob1 at cleavage site “D” to generate translation-competent 40S subunits (Fatica, Tollervey et al., 2004, Lamanna & Karbstein, 2009, Lebaron, Schneider et al., 2012). Prior to cleavage of the 20S rRNA, the pre-40S contains several AFs including Ltv1 and Enp1 bound to the pre-40S head and Tsr1, Rio2, Dim1, Pno1 and Nob1 in the vicinity of the pre-40S mRNA binding channel (Strunk, Loucks et al., 2011, Strunk, Novak et al., 2012), which are exported from the nucleus in complex with the pre-40S and released in the cytoplasm.

The release of Ltv1 and Enp1 from the pre-40S is expected to induce a major structural rearrangement during final cytoplasmic maturation that leads to the formation of a mature beak, a universal structural landmark on the head of the 40S subunit critical during translation. Beak formation also involves stable incorporation of the r-protein uS3 into the 40S and is initiated by the kinase Hrr25 in the cytoplasm (Schafer, Maco et al., 2006), which phosphorylates Ltv1 and Enp1 leading to their release from the pre-40S beak. However, the molecular basis underlying stable uS3 incorporation remains unclear, due to the lack of high-resolution structural information on pre-40S particles.

Dim1, Tsr1 and the atypical kinase/ATPase Rio2 bind to the subunit interface of the pre-40S, where they prevent the access of translation initiation factors and the 60S subunits to the immature 40S subunit (Strunk et al., 2011). Tsr1 is a GTPase-like protein, which adopts a four-domain fold similar to translational GTPases such as EF-Tu and eIF5B, but lacks the residues to bind and hydrolyze GTP (McCaughan, Jayachandran et al., 2016). Release of Rio2 is triggered by its own ATPase activity, which has been suggested to induce a conformational change in the 40S subunit (Ferreira-Cerca, Sagar et al., 2012).

Endonucleolytic cleavage of immature 20S rRNA into mature 18S rRNA by Nob1 is an essential cytoplasmic maturation event that renders the pre-40S subunit translation-competent (Fatica, Oeffinger et al., 2003, Pertschy, Schneider et al., 2009). The regulation of the rRNA cleavage involves the KH domain-containing protein Pno1 (Partner of Nob1), which co-localizes with Nob1 to the pre-40S platform. The encounter between a mature 60S subunit with a pre-40S particle is aided by the cytoplasmic GTPase Fun12 (eIF5B) and marks the point at which Pno1 is displaced and/or released and Nob1-mediated 20S pre-rRNA cleavage is triggered (Turowski, Lebaron et al., 2014). However, the molecular details of how Pno1 regulates 20S pre-rRNA cleavage remains to be clarified.

Recent cryo-EM structures of the 90S are beginning to shed light on nucleolar events of the 40S assembly (Barandun, Chaker-Margot et al., 2017, Chaker-Margot, Barandun et al., 2017, Cheng, Kellner et al., 2017, Kornprobst, Turk et al., 2016, Sun, Zhu et al., 2017). In contrast, high-resolution structural information of cytoplasmic pre-40S particles has remained inaccessible in part due to their structural flexibility and heterogeneity (Johnson, Ghalei et al., 2017, Strunk et al., 2011). We used a catalytically inactive mutant of Nob1 in budding yeast to trap and purify a cytoplasmic pre-40S particle poised for final cytoplasmic maturation. This approach permitted the structural determination of a cytoplasmic pre-40S particle at a resolution of 3.4 Å by cryo-EM. The structure provides critical insights into mechanisms that drive 40S beak formation and final pre-rRNA folding and processing and provides a mechanistic framework for the series of events that take place during pre-40S maturation.

## Results

### Isolation of cytoplasmic pre-40S poised for 20S pre-rRNA cleavage

In order to trap a pre-40S particle poised to undergo the final 20S pre-rRNA cleavage step, we exploited the catalytically inactive Nob1-D15N allele. The D15N mutation is located in the conserved PIN domain (<u>Pi</u>lT <u>N</u> terminus) of Nob1 and exhibits a dominant-negative phenotype (Fatica et al., 2004, Granneman, Petfalski et al., 2010), indicating that the nuclease activity is not required to bind pre-40S ribosomes.

We generated a conditional Nob1 mutant strain in which the endogenous *NOB1* gene was placed under the control of the GAL1 promoter (*PGAL1-NOB1*) (Appendix Table S1). On repressive glucose media, the *PGAL1-NOB1* strain was severely impaired in growth compared to *PGAL1-NOB1* complemented with plasmid-encoded Nob1 (Fig. 1a). Next, we introduced Protein A-tagged N-terminal fusions of WT Nob1 (pA-Nob1) and the catalytically inactive Nob1-D15N into the *PGAL1-NOB1* strain and analyzed growth of the resultant strains on glucose media (Fig. 1a). Consistent with previous studies (Granneman et al., 2010), the yeast cells expressing the pA-Nob1 fusion protein grew at WT rates, indicating that an N-terminal tag does not interfere with Nob1 function (Fig. 1a). In contrast, the strain expressing the catalytically inactive pA-Nob1-D15N fusion showed a severe growth defect.

**Figure 1:**
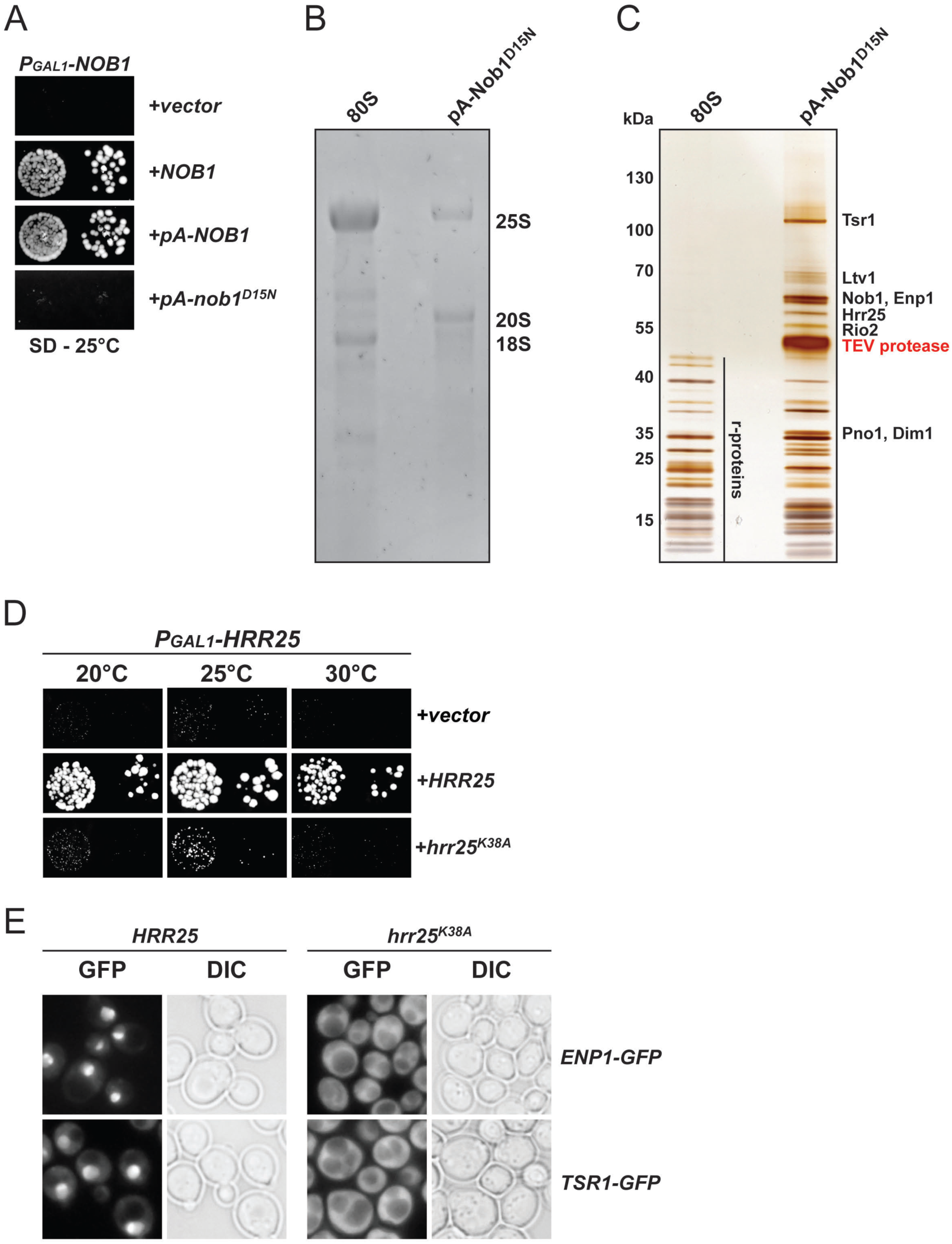
Purification of trapped pre-40S ribosomes and Hrr25 kinase activity. **A**. N-terminally tagged Protein A (pA)-Nob1 is functional, but requires endonuclease activity for viability. The *PGAL1-NOB1* strain was transformed with indicated plasmids and spotted in 10-fold dilutions on selective and repressive glucose-containing plates and grown at 25°C for 4 days. **B**. RNA composition of 80S ribosomes and purified Protein A (pA)-tagged Nob1-D15N particles. RNA was separated by agarose gel electrophoresis and stained with GelRed. **C**. Protein composition of 80S ribosomes and purified Protein A (pA)-tagged Nob1-D15N particles. Proteins were separated by SDS-PAGE and visualized by silver staining. Indicated protein bands were identified by mass spectrometry. **D**. The kinase activity of Hrr25 is essential for cell viability and is required for cytoplasmic release and recycling of Enp1 and Tsr1. Upper panel: The *PGAL1-HRR25* strain was transformed with indicated plasmids and spotted in 10-fold dilutions on selective and repressive glucose-containing plates and grown at indicated temperatures for 3-7 days. Lower panel : Yeast strains expressing endogenous Enp1-GFP or Tsr1-GFP were transformed with plasmids carrying either a galactose-inducible wild-type *HRR25* gene or a dominant-negative *hrr25*-K38A kinase dead mutant gene. Strains were then grown on galactose-containing medium at 25°C and GFP constructs were visualized by fluorescence microscopy. For a detailed description of the strains and plasmids, see Appendix Tables S1 and S2.

We affinity-purified the pA-Nob1-D15N containing ribosomal subunits from yeast cells grown in glucose-containing medium to exclusively express the mutant protein. As expected, the affinity purified particles contained 20S pre-rRNA in contrast to the mature 18S rRNA present in 80S ribosomes (Fig. 1b). In addition, we found that the isolated pre-40S particle also contained sub-stoichiometric levels of mature 25S rRNA, indicating the presence of cytoplasmic 80S-like complexes that contain mature 60S subunits (Fig. 1b). Mass spectrometric analysis of the purified sample revealed that the eight pre-40S assembly factors Hrr25, Ltv1, Enp1, Rio2, Tsr1, Dim1, Pno1 and Nob1 co-purified to nearly stoichiometric levels (Fig. 1c).

These AFs are released in a sequential manner once Ltv1 is removed (Strunk et al., 2012). In human cells, CK1δ/ε, the homologs of Hrr25, were reported to be the factors responsible for starting the cascade of AFs release (Fassio, Schofield et al., 2010, Ghalei, Schaub et al., 2015, Zemp, Wandrey et al., 2014). To investigate an analogous function of Hrr25 in yeast (Schafer et al., 2006), we generated conditional Hrr25 strains in which the endogenous *HRR25* gene or previously characterized dominant-negative mutants of Hrr25 lacking the kinase activity (Hoekstra, Liskay et al., 1991) (*hrr25*-K38A) are placed under control of a GAL1 promoter (Fig. 1d and Appendix Tables S1 and S2). Plasmids containing the galactose-inducible Hrr25 variants were transformed in yeast strains expressing endogenous GFP-tagged constructs of Enp1 or Tsr1 (Fig. 1e and Appendix Tables S1 and S2). At steady-state levels, wild-type Enp1-GFP and Tsr1-GFP localized to the nucleus. In contrast, upon overexpressing the catalytically inactive *hrr25*-K38A mutant in galactose-containing medium, both Enp1-GFP and Tsr1-GFP failed to release and recycle back to the nucleus and were therefore mislocalized to the cytoplasm. This suggests that yeast Hrr25 also acts upstream of Enp1 and Tsr1 release, as observed previously for human proteins (Zemp et al., 2014), and is likely involved in starting the cascade of AF release (Schafer et al., 2006).

### Cytoplasmic pre-40S structure determination

The affinity-purified pre-40S particles were analyzed by single-particle cryo-EM. The micrographs showed a mixture of particles that resemble 40S ribosomal subunits and 80S ribosomes in size and shape. Single-particle images were selected from the micrographs and split into subsets showing 40S-like and 80S-like particles by 2D classification. The subset of 80S-like particles was used to refine a structure at a resolution of 3.1 Å (Appendix Fig. S1 and Table S4). The 3D reconstruction revealed a mature 80S (Ben-Shem, Garreau de Loubresse et al., 2011) with additional density visible at the 3’ end of the 18S rRNA at lower local resolution, which probably represents partial density for the uncleaved 20S rRNA protruding flexibly from the 40S subunit. However, no density for any assembly factor could be observed.

The 3D reconstruction from the subset of 40S-like particles was refined reaching an overall resolution of 3.4 Å (Appendix Fig. S1 and S2 and Table S4). The density showed an immature 40S ribosomal subunit similar to the previously described cytoplasmic pre-40S ribosome (Fig. 2a) (Johnson et al., 2017, Strunk et al., 2011).

**Figure 2:**
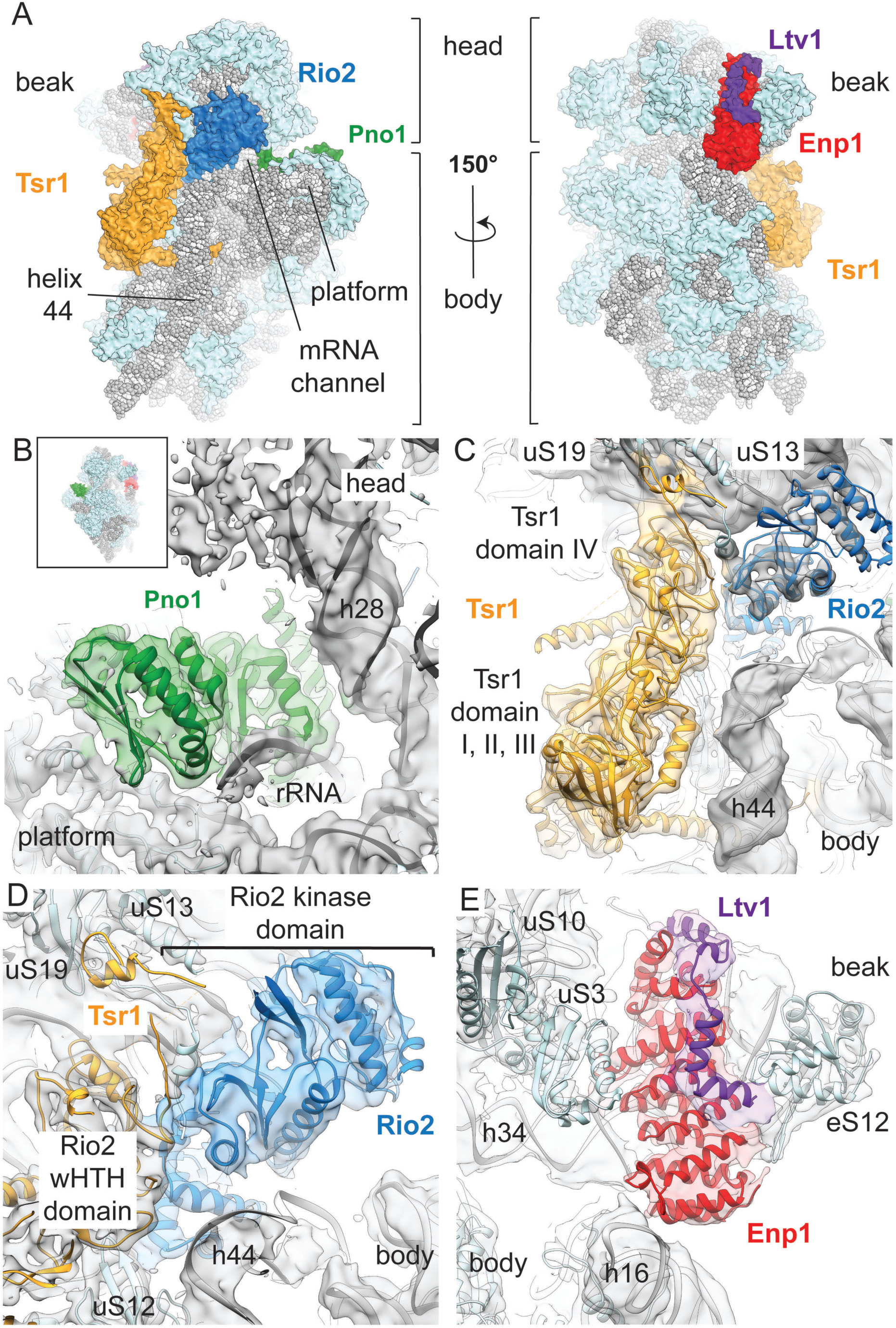
Structure of the cytoplasmic pre-40S ribosome. **A**. Overview of the front and back of the structure of the pre-40S. The 20S rRNA is depicted in gray, the r-proteins in light blue, Tsr1 in yellow, Rio2 in blue, Pno1 in green, Enp1 in red and Ltv1 in purple. **B-E.** Overview of the atomic models of the AFs in their corresponding densities. Densities were colored according to the models in a radius of 4 Å around the model atoms. For each panel, the shown densities correspond to the locally classified 3D-reconstruction used for model building. **B**. Pno1 binds to “neck” helix 28 and in vicinity of the 3’ end of the rRNA to the platform of the pre-40S particle **C**. Tsr 1 and Rio2 bind to the subunit interface and close to the decoding center (indicated with an asterisk at the top of helix 44). Tsr1 domain IV binds to the pre-40S head, while domains I-III are in contact with the body. **D**. Rio2 is also in contact with ribosomal proteins uS13 and uS19. **E**. The density of Ltv1 is shown at a different contour level (1.5σ for Ltv1 and 2.5σ for ribosome and Enp1) to highlight the visible secondary structure elements of both Enp1 and Ltv1.

The local resolution of this 3.4 Å 3D-reconstruction ranges from 2.8 Å to 4.9 Å for the ribosomal proteins and the rRNA (Appendix Figs. S2 and Table S3), allowing us to build and refine a model of the pre-40S subunit based on the 40S subunit from the 80S structure of the yeast ribosome (Ben-Shem et al., 2011) from which proteins eS10, eS26, eS31 and STM1 that are not present in our structure were removed. The reconstruction clearly revealed additional features that were assigned to the AFs by fitting previously solved crystal structures of individual yeast factors and their homologs. The distribution of the maturation factors on the surface of the pre-40S subunit forms three groups, in agreement with previously published low-resolution reconstructions of cytoplasmic pre-40S ribosomes (Johnson et al., 2017, Strunk et al., 2011). The Enp1 and Ltv1 factors are positioned between the head and the body of the subunit close to the mRNA entry channel on the solvent-exposed side of the pre-40S subunit. Tsr1 and Rio2 are localized on the side of the 40S subunit that interacts with the 60S subunit, with Tsr1 occupying the position where eIF5B binds during translation initiation as previously reported (Johnson et al., 2017, McCaughan et al., 2016, Strunk et al., 2011), while Rio2 is positioned in the middle of the mRNA channel bridging the head and the body of the 40S subunit. Pno1 binds on the soluble side of the subunit behind the platform where the 3’ end of the 18S rRNA is located in the mature 40S subunit. The average local resolution of the densities corresponding to Tsr1 (3.9 Å), Pno1 (3.7 Å), and Rio2 (4.0 Å) allowed us to rebuild their structures in complex with the ribosome, whereas the density corresponding to Enp1 and Ltv1 was resolved at lower local resolution (4.3 Å) and was interpreted by rigid-body fitting the crystal structure of the partial Enp1-Ltv1 complex from yeast (PDB : 5WWO (Sun et al., 2017)) (Fig. 2b-e and Appendix Figs. S2 and Table S3).

The Dim1 rRNA dimethylase reported to be part of the pre-40S could only be identified in a small fraction of the final set of 40S-like particles after additional local classification (cf. Appendix Figs. S1 and S3 and Table S4). The reconstruction of the pre-40S subunits containing Dim1 shows helix 45 in its mature position and Dim1 at a position too distant from its target helix 45 for methylation (Boehringer, O’Farrell et al., 2012, Johnson et al., 2017) (Appendix Fig. S3). This suggests that Dim1 might have already methylated h45 and remains only weakly attached to the rRNA.

### Configuration of Enp1, Ltv1 and uS3 on the pre-40S head

On the solvent-exposed side of the pre-40S ribosome, densities corresponding to Enp1 and Ltv1 connect the beak of the pre-40S head with rRNA helix 16 on the pre-40S body (Fig. 2a and 2e). The superhelical fold of Enp1 fits well into the density and an additional L-shaped density on top of Enp1 could be attributed to residues 362-405 of Ltv1 in agreement with the high-resolution Enp1-Ltv1 crystal structure (Fig. 2e) (Sun et al., 2017). The large positively charged surface of Enp1 contacts the three-way junction of rRNA helices 32, 33 and 34 in the beak (Appendix Fig. S4), where eS10 would bind in the mature 40S (Ben-Shem et al., 2011), and is likely stabilizing these helices in an immature conformation (Fig. 3a). The three helices are highly distorted compared to the mature 40S, changing the overall shape of the beak.

**Figure 3:**
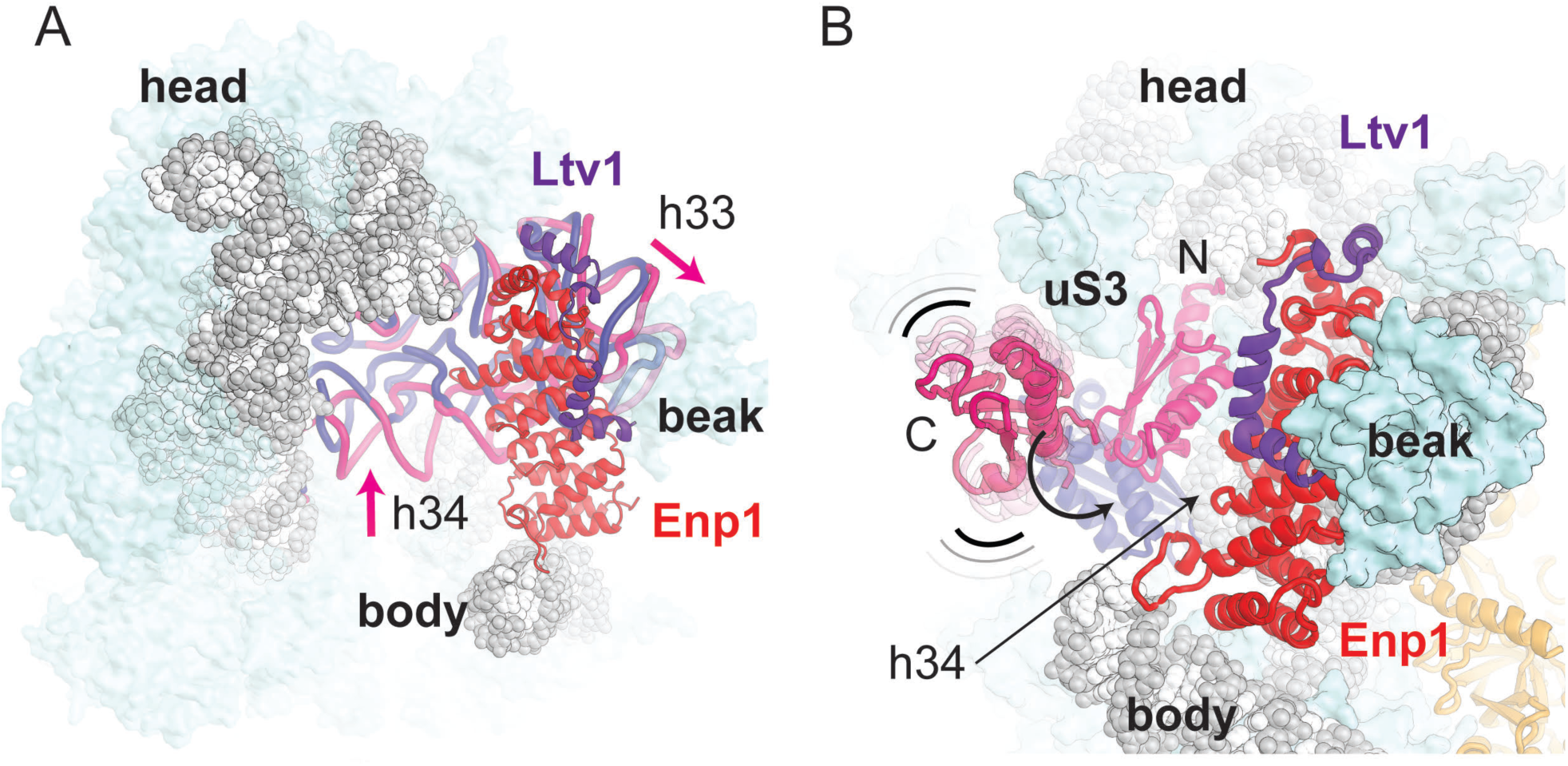
Enp1 and Ltv1 interact with the distorted rRNA in the head of the pre-40S. **A**. Overview of the head rRNA with the r-proteins shown as light-blue surfaces, Enp1 and Ltv1 as cartoon, and rRNA as gray spheres and cartoon. uS10, uS3 and uS14 were removed for clarity. The rRNA corresponding to helix 32, 33, and 34 is shown as cartoon to highlight the change in conformation between our model (pink) and the mature 40S rRNA (dark blue, aligned to helix 42 and 43 of our model). **B**. Position of the flexibly bound uS3 C-terminal domain in our structure (pink) and the stably incorporated uS3 C-terminal domain in the context of the mature 40S subunit (dark blue). Within the pre-40S particle, the uS3 C-terminal domain cannot adopt its mature position, as the displaced helix 34 is in its way.

Enp1 contacts the ribosomal protein uS3 N-terminal domain, which occupies its mature position with respect to helix 41 and uS10 (Fig. 3b). On the other hand, the uS3 C-terminal domain does not localize to its mature position, which is occupied by the displaced h34. It is only visible at low contour level as a density disconnected from the core of the pre-40S and hence this domain is likely to be integrated once h34 withdraws to its final conformation (Fig. 3b). This provides a molecular explanation for the previously observed lower affinity of uS3 to pre-40S particles (Schafer et al., 2006). Moreover this finding implies that the Hrr25-dependent stable incorporation of uS3 into the pre-40S is linked to a large-scale conformational re-arrangement of rRNA in the beak upon departure of Enp1.

### Tsr1 and Rio2 contact multiple functionally important sites on the 40S

The crystal structure of Tsr1 (PDB 5IW7 (McCaughan et al., 2016)) could be fitted and refined into an extra density located between the head and the body of the cytoplasmic pre-40S, with only minimal adjustments of domain IV (Fig. 2c). Tsr1 binds on top of uS12 (Fig. 4a), a highly conserved ribosomal protein linked to translational accuracy (Alksne, Anthony et al., 1993, Triman, 2007). At the ribosomal P-site close to domain IV of Tsr1 we observe density for Rio2, which could be interpreted by separately fitting homologous crystal structures of the N- and C-terminal halves of Rio2 from *C. thermophilum* (Ferreira-Cerca et al., 2012) (Fig. 2d). Interestingly, the eukaryote-specific C-terminal extension segment that contains the autoinhibitory helix (Ferreira-Cerca et al., 2012) is disordered in our structure, as expected for Rio2 in an active conformation. Together, Tsr1 and Rio2 are blocking the A- and P-site on the 40S subunit where tRNAs and initiation factors bind.

**Figure 4:**
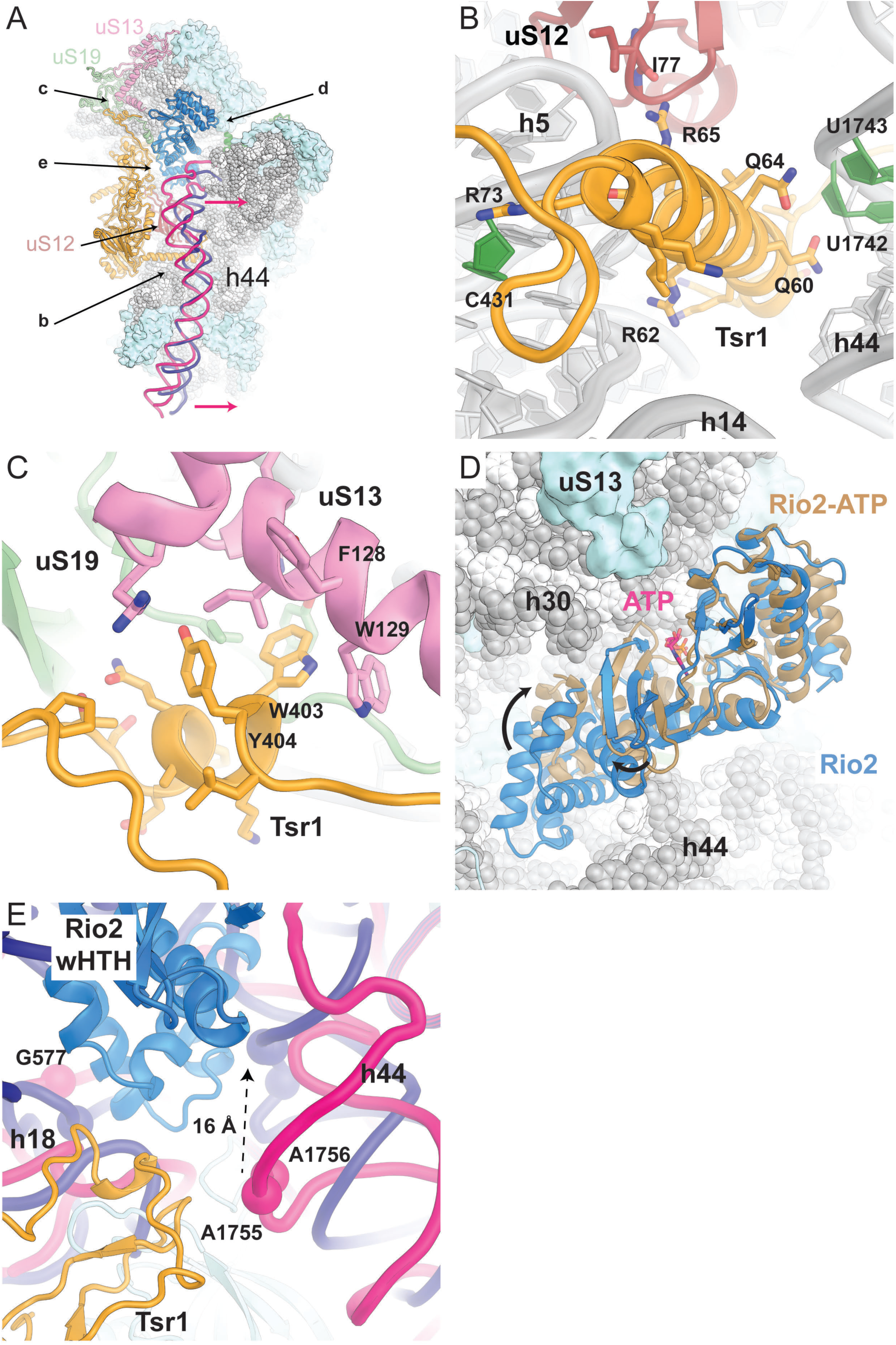
Tsr1 and Rio2 interact with functionally important regions of the 40S subunit. **A**. Overview of the pre-40S ribosome. The rRNA is represented as spheres (gray) and the r-proteins as surfaces (light blue). Rio2 (blue), Tsr1 (orange), uS12 (red), uS13 (pink), and uS19 (light green) are represented as cartoon. To help with orientation arrows indicate the viewing perspectives used to generate panels b-e. The pre-40S helix 44 (cartoon in pink) is highly distorted compared to the mature one (dark blue, PDB 4V88 (Ben-Shem et al., 2011)). **B**. The conserved N-terminal α-helix of Tsr1 (residues 51-76) (Appendix Fig. S6) is wedged between the body of the pre-40S and helix 44 causing its displacement (Panel A). **C**. The highly conserved stretch of Tsr1 residues 391-406 (Appendix Fig. S6) interacts with uS13 and uS19. Tsr1 W403 is inserted into a hydrophobic pocket formed by uS13 F128, W129 and Tsr1 Y404. **D**. Comparison of the yeast pre-40S Rio2 conformation (blue) and the crystal structure of ATP-bound Rio2 from *A. fulgidus* (brown, PDB : 1ZAO, (LaRonde-LeBlanc et al., 2005)). The structures are superimposed relative to their kinase domains. The ATP visualized in the crystal structure is shown in pink to highlight the binding pocket. **E**. Close-up view of the decoding center, highlighting the shift in the decoding bases A1755 and A1756 between the Pre-40S particle (pink) and the mature ribosome (dark blue, PDB 4V88 (Ben-Shem et al., 2011))

Remarkably, the Tsr1 density forms a helical structure that inserts between the body of the small subunit and helix 44 (Fig. 4b and Appendix Fig. S5). This α-helix corresponds to residues 51-75 of Tsr1 that were not resolved in the previously reported Tsr1 crystal structure (McCaughan et al., 2016). The conserved positively charged amino acids (Appendix Fig. S6) of this Tsr1 α-helix directly interact with the minor groove of rRNA helix 44, as well as with rRNA helices 5, 11 and 24, and protein uS12 (Fig. 4b and Appendix Fig. S5).

In our cytoplasmic pre-40S reconstruction rRNA helix 44 is considerably displaced compared to its position in the mature 40S (Fig. 4a) and its conformation would be incompatible with the joining of the 60S subunit. The conformational rearrangements also propagate to the top of helix 44, which forms the ribosomal A-site, and result in a displacement of the decoding center bases 1755 and 1756 by approximately 16 Å (Fig. 4e and Appendix Fig. S5). This displacement indicates a possible role for Tsr1 in preventing the premature formation of the decoding center. Additionally, the shift of helix 44 at the decoding center allows the insertion of Rio2 (Fig 4e).

Furthermore, we could assign extra densities to residues 380 to 409 of Tsr1 belonging to a long linker between Tsr1 domains II and domain III, which had remained unresolved in the Tsr1 crystal structure due to conformational flexibility. The linker first contacts Rio2 and the C-terminus of uS19 and extends along Tsr1 domain IV in the direction of the pre-40S head. The last visible residues of the linker form specific contacts with the displaced rRNA helix 31 as well as with proteins uS13 and uS19 (Fig. 4c and Appendix Fig. S5). These residues, 392 to 408, are highly conserved in eukaryotes (Appendix Fig. S6), pointing towards a key role of this interaction with these two universally conserved ribosomal proteins, whose prokaryotic homologs are important for intersubunit bridging (Bowen, Musalgaonkar et al., 2015, Jenner, Demeshkina et al., 2010) as well as for tRNA binding (Graifer & Karpova, 2015, Lomakin & Steitz, 2013).

Rio2 is bound at the decoding center of the 40S (Fig. 4e). The Rio2 N-terminal winged-helix-turn-helix (wHTH) domain is inserted between 20S rRNA helices 1, 18, 28, 34 and 44, which form the decoding center of the mature 40S, while the bi-lobed C-terminal kinase domain blocks the cytoplasmic pre-40S P-site from tRNA binding (Fig. 4a). The wHTH domain interacts with and distorts helices 18 and 44, restraining their conformation and preventing premature folding of the decoding center (Fig. 4e).

Interestingly, the structure of cytoplasmic pre-40S-bound Rio2 shows that the two halves of the Rio2 kinase domain, which form the nucleotide binding pocket, are rotated relative to each other when compared to the crystal structures of the *A. fulgidus* (LaRonde-LeBlanc, Guszczynski et al., 2005, LaRonde-LeBlanc & Wlodawer, 2004) (Fig. 4e) and *C. thermophilum* homologs (Ferreira-Cerca et al., 2012). As a result, the residues responsible for ATP binding are separated by greater distances, which most likely renders Rio2 unable to cleave ATP.

### Pno1 interacts with the 3’ end of rRNA at the pre-40S platform

Pno1 is the interaction partner of the endonuclease Nob1 that mediates the cleavage of 20S rRNA into mature 18S rRNA (Woolls, Lamanna et al., 2011). On the platform, close to the 40S “neck”, where eS26 is located in the mature 40S, additional density could be assigned to Pno1 (Fig 2b). The Pno1 KH-like and KH domains were modeled using the crystal structure of the homologous *P. horikoshii* Pno1 protein (Jia, Horita et al., 2010). We observed that the C-terminal part of Pno1 would clash with the mature position of eS28 and helix 28 (Fig 5a), indicating that the tilted conformation of the cytoplasmic pre-40S head is required for Pno1 binding. Interestingly, the KH domain is also interacting with the loop between helices 44 and 45, which is in a stretched-out conformation due to the displaced position of helix 44 in the cytoplasmic pre-40S.

**Figure 5:**
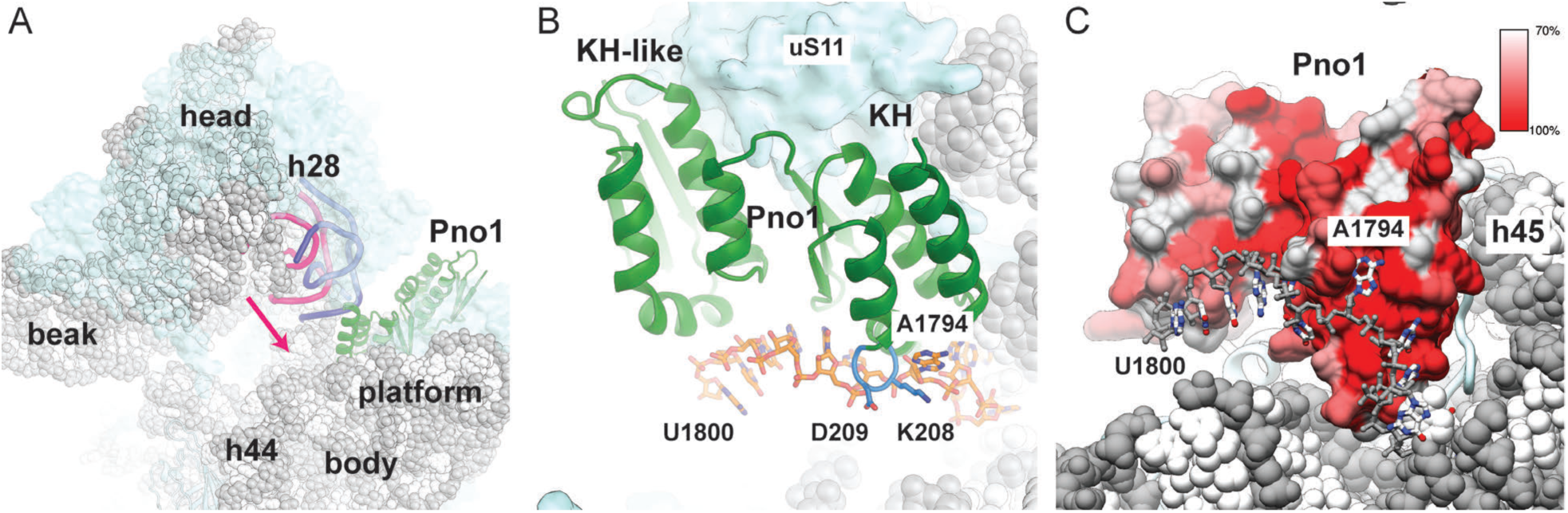
Pno1 interacts with the rRNA of the platform. **A**. Overview of the pre-40S ribosome. The rRNA is represented as spheres (gray) and the r-proteins as surfaces (light blue). In the pre-40S particle, helix 28 (cartoon in pink) is highly distorted compared to the mature conformation (dark blue, PDB 4V88 (Ben-Shem et al., 2011)) in which it would clash with Pno1 (green). **B**. Pno1 interacts with 20S rRNA nucleotides 1787-1800. Nucleotide A1794 is specifically recognized by the C-terminal KH domain of Pno1, which contains a conserved GxxG RNA binding motif (blue). A1794 is flipped out, which twists the RNA such that the bases that follow bind to the surface of Pno1. **c.** Surface representation of Pno1 colored by residue conservation. Highly conserved residues (red) form the surface interacting with the 20S rRNA (Appendix Fig. S7).

Our density reveals that Pno1 binds the 3’ end of the rRNA at the location that would precede the final D cleavage site at nucleotide 1800 targeted by Nob1 (Fig 5b and Appendix Fig. S5). Particularly, nucleotides 1793 to 1795 are recognized in a flipped out conformation by specific interactions with the highly conserved GXXG motif of the C-terminal KH domain of Pno1, whereas the RNA up to the cleavage site is twisted such that the bases point away from the surface of the protein (Fig. 5c, Appendix Fig. S5) and fades for the remainder of 20S pre-rRNA sequence into featureless density.

Despite being used as the bait to isolate the pre-40S particle, no density could be assigned to Nob1-D15N at the D cleavage site where Nob1 was reported to be located (Johnson et al., 2017, Strunk et al., 2011) even after further focused classification (Scheres, 2012). However, it is possible that Nob1 is flexibly bound to downstream parts of the 20S rRNA (Lamanna & Karbstein, 2009, Turowski et al., 2014).

## Discussion

The reconstruction of the cytoplasmic pre-40S subunit at 3.4 Å resolution revealed how the AFs check the integrity of functionally important regions of the subunit and lock the pre-rRNA conformation to prevent premature progression through the maturation process and prepare the pre-rRNA for processing. We observe striking differences between the cytoplasmic pre-40S state and the mature 40S ribosome exemplified by the distortion of the 3’ minor and 3’ major domains of the rRNA that is accompanied by an extreme rotation of the cytoplasmic pre-40S head that exceeds any 40S head rotation observed during translation. This stage of maturation seems to be focused on the 3’ major domain of the rRNA since it is contacted by all observed AFs except Pno1.

### Final maturation of the pre-40S beak

In our structure, the large positively charged surface of Enp1 directly interacts with the beak rRNA, most likely acting as a scaffold and restraining it in its observed distorted conformation. The protein kinase Hrr25 was reported to regulate the cytoplasmic maturation of the pre-40S through the phosphorylation of Enp1 and Ltv1, leading to the release of Ltv1 (Ghalei et al., 2015), and stable incorporation of uS3 (Schafer et al., 2006). Since the displaced helix 34 prevents the formation of the mature binding pocket for the uS3 C-terminal domain, Hrr25-dependent incorporation of uS3 must be linked to the conformational rearrangement of helix 34. One possibility is that the release of Ltv1 after phosphorylation of Ltv1 and Enp1 by Hrr25 (Ghalei et al., 2015) destabilizes the binding of Enp1, which dissociates and allows the beak and uS3 to adopt their mature conformations and the head to rotate which in turn changes the conformation of Rio2 (Fig. 6).

**Figure 6:**
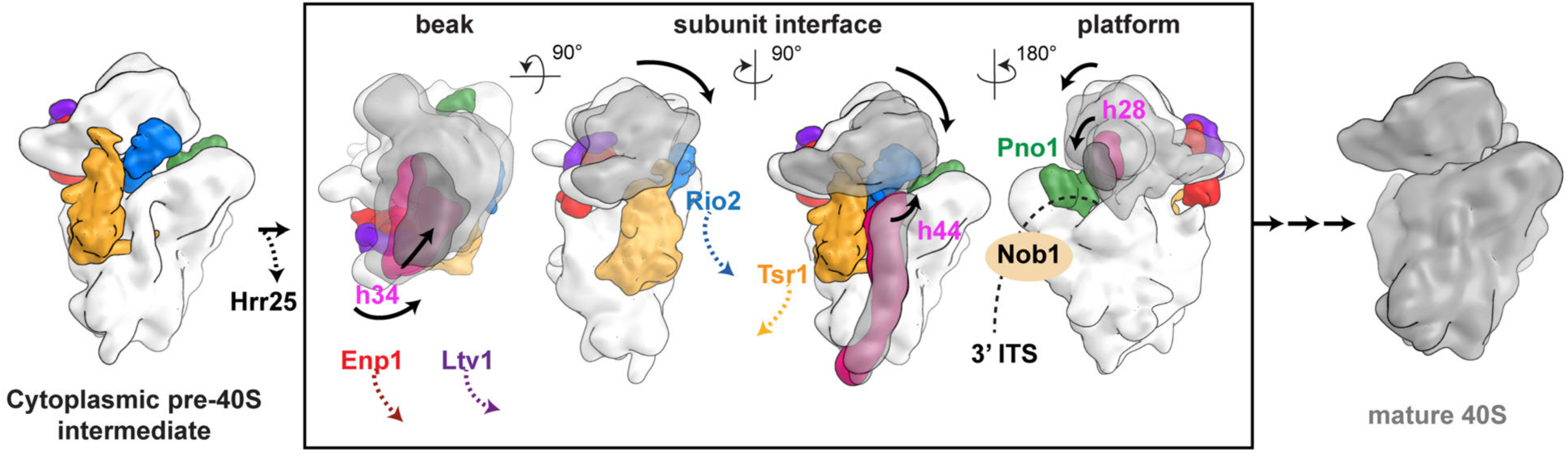
A model for cytoplasmic maturation of the pre-40S subunit through series of interdependent conformational changes. The boxed region of the scheme shows the pairwise changes in rRNA conformation between the pre-40S (white) and the mature 40S (gray) that are linked to the release of assembly factors without implying a sequential order. The tilt and rotation of the head as well as conformational changes in helix 34, helix 44 and helix 28 (pink) are interconnected such that conformational changes in the beak, the subunit interface and the platform lead to the release of AFs from their respective binding sites.

### Tsr1 and Rio2 probe the inter-subunit interface and the decoding center

It has been suggested that the role of Tsr1 is to prevent the premature binding of 60S ribosomal subunit by sterically blocking the access of the 60S subunit (Johnson et al., 2017, McCaughan et al., 2016, Strunk et al., 2011). In the structure described here, we additionally observe that Tsr1 is responsible for stabilizing helix 44 in a shifted conformation compared to the mature 40S, which also clearly prevents the premature binding of the 60S.

The two domains of Rio2 are bound to the head and the body of the subunit and therefore their relative positioning is linked to the head rotation. In the observed conformation, Rio2 cannot be active in ATP hydrolysis since the two lobes of the kinase domain forming the nucleotide binding cleft are rotated away from each other compared to the ATP-or ADP-bound structures of Rio2 homologs *from A. fulgidus* (LaRonde-LeBlanc & Wlodawer, 2004)(LaRonde-LeBlanc et al., 2005). Consequently, head rRNA rearrangement to its mature conformation upon dissociation of Enp1-Ltv1 could possibly induce a closure of the Rio2 nucleotide binding cleft, thereby activating Rio2 to hydrolyze ATP that leads to autophosphorylation and to dissociation (Fig. 6) (LaRonde-LeBlanc et al., 2005) (Ferreira-Cerca et al., 2012). Since both Rio2 and Tsr1 interact with helix 44 of the pre-rRNA and they furthermore contact each other (Fig. 4a), dissociation of Rio2 following autophosphorylation would lead to destabilization and dissociation of Tsr1. This allows the formation of the decoding center and accommodation of rRNA helix 44 to its mature conformation propagating conformational changes towards the platform where Pno1 interacts with the 3’ end of the rRNA.

The contacts of Rio2 and Tsr1 with the decoding center and important regions around it, including RNA helices 18 and 44, and proteins uS12, uS13, and uS19, point towards an additional function for Rio2 and Tsr1 as AFs that monitor the proper formation of this key functional region of the 40S subunit. As Rio2 and Tsr1 bind the pre-40S in the nucleus and are needed for transport into the cytoplasm (Schafer, Strauss et al., 2003), these interactions could serve as one of the “checkpoints” for nuclear export by preferentially binding only to pre-40S particles, which contain these r-proteins.

### Repositioning of Pno1 is required for 3’ end processing

Our structure also reveals how Pno1 is bound to the 20S RNA in the region corresponding to the 3’-end of the mature rRNA in the immediate vicinity of site D where Nob1 endonuclease binds (Turowski et al., 2014). In this conformation, the close proximity of Pno1 to adjacent rRNA elements would prevent access of the Nob1 PIN domain to the D cleavage site, which explains why pre-rRNA processing does not take place at this stage of maturation. Considering that Pno1 occupies the binding site for eS26 protein, which is not visible in our pre-40S structure, it is possible that eS26 insertion, coupled with a repositioning of Pno1, is required for cleavage of the 20S pre-rRNA by Nob1. Notably, provided that Pno1 moves from its current location, Nob1 would be able to interact with the KH-like domain of Pno1, as observed biochemically (Woolls et al., 2011), and cleave the rRNA. In support of this notion, it was recently shown that a conformational change in Pno1 is required to allow Nob1 access for 20S pre-rRNA cleavage (Turowski et al., 2014, Widmann, Wandrey et al., 2012) and that eS26-depletion impairs final cytoplasmic 20S pre-rRNA processing (Schutz, Fischer et al., 2014).

### Interdependent AF departure events lead to 40S maturation

The structure of the pre-40S subunit described here reveals the molecular basis of concerted events that take place upon departure of the visualized AF. Although it is not possible to suggest the exact temporal sequence of maturation events, we now understand the interdependence of various AFs in stabilizing a particular conformation of the pre-rRNA (Fig. 6). Based on the structure and the available biochemical and genetic evidence, the process is likely to start with the phosphorylation of Ltv1 and Enp1 by Hrr25 that destabilizes their interactions with the beak of the pre-40S subunit. This leads to a conformational change in the rRNA allowing accommodation of the C-terminal domain of uS3. After or during beak formation, the 40S head likely rotates to its mature position, triggering ATP hydrolysis by Rio2 and its dissociation. Since both Rio2 and Tsr1 stabilize helix 44 in a distorted conformation, dissociation of Rio2 will also lead to dissociation of Tsr1. Accommodation of helix 44 into its mature position will lead to conformational changes in the rRNA at the platform, allowing formation of rRNA helix 28 and docking of eS26, which then repositions Pno1 and allows binding of Nob1 for pre-rRNA cleavage. This mechanism also implies that Nob1 would only have access to Pno1 once the 40S matures to the point of being able to associate with the 60S subunit, in agreement with previous biochemical results (Fig 6) (Lebaron et al., 2012, Strunk et al., 2012, Turowski et al., 2014).

In conclusion, our structure sheds new light on the cytoplasmic maturation pathway of the 40S subunit by revealing contact sites of the assembly factors Ltv1, Enp1, Rio2, Tsr1 and Pno1, and their interactions with 20S rRNA. The cytoplasmic pre-40S exhibits a highly distorted 3’ major and 3’ minor rRNA domain conformation stabilized by the AFs, which suggests a mechanism for the concerted release of the AFs orchestrated by the folding of the rRNA in the head of the pre-40S subunit during the final stages of maturation. These results provide the structural basis for biochemical and genetic experiments to dissect the effects of the observed interactions on the cascade of mechanistic events leading to the formation of the mature 40S subunit.

## Materials & Methods

### Purification of pre-40S ribosomes

Pre-40S ribosomes were purified from a *Saccharomyces cerevisiae* strain expressing the catalytically inactive Nob1-D15N mutant fused to an N-terminal and cleavable Protein A tag (pA-TEV-FLAG-Nob1D15N). For this purpose, a *NOB1* conditional mutant strain (*P_Gal1_-NOB1*) was transformed with a plasmid encoding the tagged Nob1 mutant gene (pRS315-pA-TEV-FLAG-Nob1D15N). Cells were grown at 25°C in glucose-containing synthetic medium to an optical density of 1.5 and then harvested at 4000 x g at 4°C for 15 min. Pellets were first washed with ddH_2_O and then with one pellet volume of 20 mM HEPES pH7.4 buffer supplemented with 1.2% polyvinylpyrrolidone and protease inhibitors (Roche Complete, PMSF). The final pellet was transferred to a 20 ml syringe and pushed directly into liquid nitrogen to produce frozen yeast noodles. Cell lysis was performed by cryo-grinding in a Planetary Ball Mill (Retsch Pulverisette 6) to generate fine yeast powder to be stored at −80°C. To purify pre-40S ribosomes, 3-5 g of yeast powder was resuspended in 50 ml of buffer A (20 mM HEPES pH7.4, 110 mM KOAc, 40 mM NaCl, 10 mM MgCl_2_, 0.5% Triton-X100, 0.1% Tween-20, PMSF), cleared by centrifugation at 4000 x g for 5 min and then filtered with a syringe first through a 2.7 µm and then through a 1.6 µm filter (Whatman). The cleared lysate was incubated on a nutator for 30 min at 4°C with a 100 µl slurry of magnetic Dynabeads (Epoxy M-270, Thermo Fischer Scientific) conjugated with rabbit IgG antibodies (Sigma Aldrich). IgG antibodies were conjugated to Dynabeads as described (Oeffinger, Wei et al., 2007). After incubation, beads were collected with a magnet rack and washed five times in 1 ml of buffer A and then three times in 1 ml of buffer B (buffer A without Triton-X100 and Tween-20). Finally, beads were resuspended in 50 µl of buffer B and then incubated at 4°C overnight with 10 units of TEV protease (Sigma Aldrich) to elute the purified pre-40S ribosome. Mature 80S ribosomes were purified from *Saccharomyces cerevisiae* as described previously (Greber et al., 2016) and using associative conditions (20 mM HEPES pH7.4, 100 mM KCl, 20 mM MgCl_2_) in the final gradient.

The RNA composition of ribosomal particles was determined by gel electrophoresis (2% w/v agarose) followed by GelRed (Biotium) staining, while the protein composition was determined by SDS-PAGE (NuPAGE 4-12% v/v Bis-Tris, Thermo Fischer Scientific) followed by silver staining. For protein identification, silver-stained bands were cut out and identified by mass spectrometry.

### Yeast methods and fluorescence microscopy

The *Saccharomyces cerevisiae* strains used in this study are listed in Appendix Table S1. Genomic disruptions, C-terminal tagging and promoter switches at genomic loci were performed as described previously (Janke, Magiera et al., 2004).

Plasmids used in this study are listed in Appendix Table S2. Mutant constructs of *NOB1*, *HRR25*, *RIO2* and *TSR1* were generated using the QuikChange site-directed mutagenesis kit (Agilent Technologies). All cloned DNA fragments and mutagenized plasmids were verified by sequencing.

To monitor subcellular localization of Enp1 and Tsr1, yeast cells expressing catalytically inactive variants of Hrr25 (Hrr25-K38A) and GFP-tagged Enp1 or Tsr1 constructs were grown in YPD at 25°C to log phase. Cells were visualized using a DM6000B microscope (Leica Microsystems) equipped with a HCX PL Fluotar 63×/1.25 NA oil immersion objective (Leica Microsystems). Images were acquired with a fitted digital camera (ORCA-ER; Hamamatsu Photonics) and Openlab software (Perkin-Elmer).

### EM sample preparation and data collection

Quantifoil R2/2 holey carbon copper grids (Quantifoil Micro Tools) were glow-discharged for 15 seconds at 15 mA using a easiGlow Discharge cleaning system (PELCO) prior to coating with graphene oxide (Pantelic, Meyer et al., 2010). The sample at 70 ng/µL was used for cryo-grid preparation using a Vitrobot (FEI Company), whose chamber was set to 4 ˚C and 100% humidity. After incubating for 90 seconds in the chamber with 5 µL of sample, the grids were blotted for 12-15 seconds and rapidly plunge-frozen in a mixture 2:1 of Propane:Ethane (Carbagas) at liquid nitrogen temperature.

One grid was used for data collection in a Titan Krios cryo-transmission electron microscope (FEI Company) operating at 300 kV and equipped with a Falcon 3EC. A dataset of 3083 micrographs was collected using the EPU software (FEI Company) in a range of defocus between −0.6 and −2.5 µm and a magnification of 100720 (pixel size of 1.39 Å/pixel). Each micrograph was collected in integration mode split into 38 frames using a total dose of 40 electrons/Å^2^.

### Data processing and reconstruction

The stacks of frames were aligned to correct for beam-induced motion and dose-weighted using MotionCor2 (Zheng, Palovcak et al., 2017). The non-dose-weighted micrographs were used to estimate the Contrast Transfer Function (CTF) using GCTF (Zhang, 2016). 2867 good micrographs were selected by evaluating the quality of their CTF estimation, the quality of their power spectra, and the quality of the micrographs themselves. From these, roughly 595000 particles were picked using GAutomatch (http://www.mrc-lmb.cam.ac.uk/kzhang/) using as template 2D averages of 5000 particles picked without reference. These particles were then processed using Relion 2.1 (Kimanius, Forsberg et al., 2016). The particles were extracted and binned 5 times (pixel size of 6.95 Å/pixel) before being used for a reference-free 2D classification. After removing images representing ice crystals and carbon edges as well as 80S particles, approximately 236000 particles were selected for refinement using a mature 40S density as reference followed by a 3D classification without alignment of the particles. The 165000 good particles containing Tsr1 were selected for a refinement using the full pixel size (1.39 Å/pixel). After post-processing and sharpening, the reconstruction reached an overall resolution of 3.4 Å (Fig. S2).

Further focused classifications were performed with different masks around densities corresponding to ENP1, Rio2, Dim1, and Pno1 generated manually using UCSF Chimera (Pettersen, Goddard et al., 2004).

### Modeling and Docking

First, atomic models of the 40S head and body obtained from the crystal structure of the mature *S. cerevisiae* 80S (PDB : 4V88 (Ben-Shem et al., 2011)) were docked as rigid bodies into the EM density of the high-resolution 40S after removal of eS10, eS26, eS31 and STM1, which are not visible from the pre-40S, using UCSF Chimera (Pettersen et al., 2004). uS3 was split into its N- and C-terminal domains, and each domain was docked independently. For initial interpretation of the AFs, the crystal structures of yeast Tsr1 (PDB : 5IW7, (McCaughan et al., 2016)) and the yeast Enp1-Ltv1 complex (PDB : 5WWO, (Sun et al., 2017)) as well as homology models based on the crystal structures of *A. fulgidus* Rio2 (PDB : 1ZAO, (LaRonde-LeBlanc et al., 2005)), human Dim1 (PDB : 1ZQ9) and *Pyrococcus horikoshii* Pno1 (PDB: 3AEV, (Jia et al., 2010)), which were generated using PHYRE2 (Kelley, Mezulis et al., 2015), could be unambiguously fitted into the EM densities obtained from the focused alignment runs (Fig. 2 and Appendix Fig. S1). Due to the limited resolution obtained for the densities of Dim1 and the Enp1-Ltv1 complex, these AFs were docked as rigid bodies, while the remaining AFs were resolved between 3.6 – 4.0 Å (Appendix Table S3), which allowed manual rebuilding using COOT (Emsley, Lohkamp et al., 2010) and O (Jones, 2004). In the r-proteins, major structural differences between the mature 40S and the pre-40S were observed in the vicinity of the AFs, and these areas were readjusted similarly. The rRNA was rebuilt by docking of segments as rigid bodies and rebuilding using COOT. Major rRNA rearrangements included the beak and helix 34 in the pre-40S head as well as helix 44 in the body. Additional structural deviations between the 40S crystal structure and the EM reconstruction were corrected in areas with sufficient resolution. With the exception of Dim1, a final pre-40S model was then assembled from the coordinates built into the individual EM densities obtained after focused alignment and fitted into the high-resolution EM map. To regularize the geometry and optimize the fit of the final atomic model to cryo-EM map, the structure was then refined in reciprocal space using PHENIX.REFINE (Adams, Afonine et al., 2010) against structure factors back-calculated from the high-resolution EM density as described earlier (Greber, Boehringer et al., 2014) (Appendix Table S4). The refinement was stabilized in areas of weaker density by applying Ramachandran restraints and secondary structure restraints for protein α-helices and β-strands as well as rRNA base pairs. An optimal weighting of the model geometry versus the structure factors during refinement was obtained with a wxc value of 1.4. The model was validated by comparison of the FSCs calculated for the experimental density and the model (Appendix Fig. S2).

### Figure generation

Local resolutions were estimated using the “relion_postprocess” from Relion2.1 (Kimanius et al., 2016). Figures showing cryo-EM structures were generated using UCSF Chimera (Pettersen et al., 2004) and Pymol (The Pymol Molecular Graphics System Version 1.8.4.0 Schrödinger). The electrostatic surface charge distribution was calculated using the APBS (Adaptive Poisson-Boltzmann Solver) (Baker, Sept et al., 2001, Lerner, 2006) plug-in of Pymol. Multisequence alignments were performed using Clustal Omega (Sievers, Wilm et al., 2011) and visualized with ESPript (Robert & Gouet, 2014).

## Acknowledgements

We would like to thank the ETH Zürich scientific centre for optical and electron microscopy (ScopeM) for access to electron microscopy equipment and are indebted to P. Tittmann for technical support. We thank all members of the Panse laboratory for enthusiastic discussions, and the Institute of Medical Microbiology and the UZH for continued support.

VG Panse is supported by grants from the Swiss National Science Foundation, NCCR RNA & Disease, Novartis Foundation, Olga Mayenfisch Stiftung and a Starting Grant Award from the European Research Council (EURIBIO260676). This work was supported by the Swiss National Science Foundation grant and via the National Centre of Excellence in RNA and Disease & project funding 138262 to N. Ban.

## Author contributions

C.P. and V.G.P. conceived the project. C.P. designed the system to isolate the trapped pre-40S, performed the purification and validated the preparations by negative stain electron microscopy. A.S. prepared grids for cryo-EM, collected electron micrographs and processed them with the aid of D.B. The atomic model was interpreted by M.L. and M.W. S.G. and P. K-N. performed yeast *in vivo* experiments. All authors were involved in the interpretation of results. A.S., C.P, M.W., D.B., V.G.P and N.B. wrote the manuscript.

## Data and materials availability

The final model containing the cytoplasmic Pre-40S with Enp1, Ltv1, Tsr1, Rio2, and Pno1 is available from the Protein Data Bank (PDB) under accession number XXXX. The cryo-EM map of the unclassified pre-40S is deposited in the Electron Microscopy Data Bank (EMDB) under accession number EMD-XXXX, as well as the reconstructions from the focus classifications, which can be found under the accession numbers EMD-XXX (Rio1), EMD-XXXX (Enp1), and EMD-XXXX (Dim1), respectively. The map of the empty 80S is deposited in the EMDB under accession number EMD-XXXX.

## Appendix information for Structure of the eukaryotic cytoplasmic pre-40S ribosomal subunit

**Appendix Figure S1:**
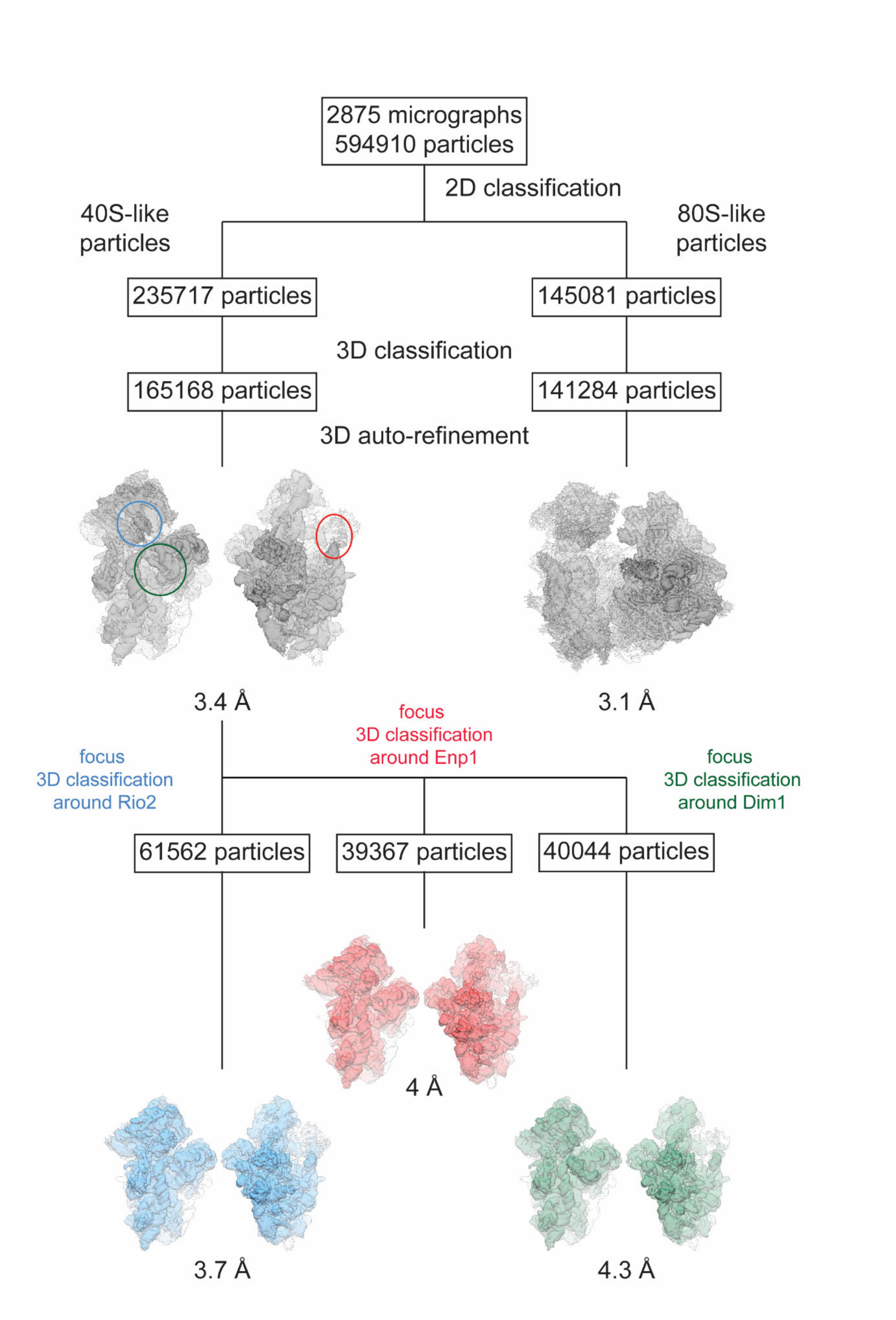
Classification and refinement scheme for the Pre-40S complex, the 80S-like particle and the Dim1-containing complex. 2D-classification from 594,910 particles that had been picked from 2,875 micrographs revealed two particle subpopulations of “40S-like” and “80S-like” particles, which where separately processed and subjected to 3D-classification and auto-refinement yielding a 3D-reconstruction of the Pre-40S complex at an overall resolution of 3.4 Å and a 3D-reconstruction of the 80S-like particle at an overall resolution of 3.1 Å. Subsequent rounds of focused 3D-classification from this final set of Pre-40S particles focussing on either Rio2 (blue circle), Enp1 (red circle) or Dim1 (green circle) yielded subpopulations of the Pre-40S complex at overall lower resolution but with higher local resolution in the area of focus, further facilitating the model building and docking in these areas.

**Appendix Figure S2:**
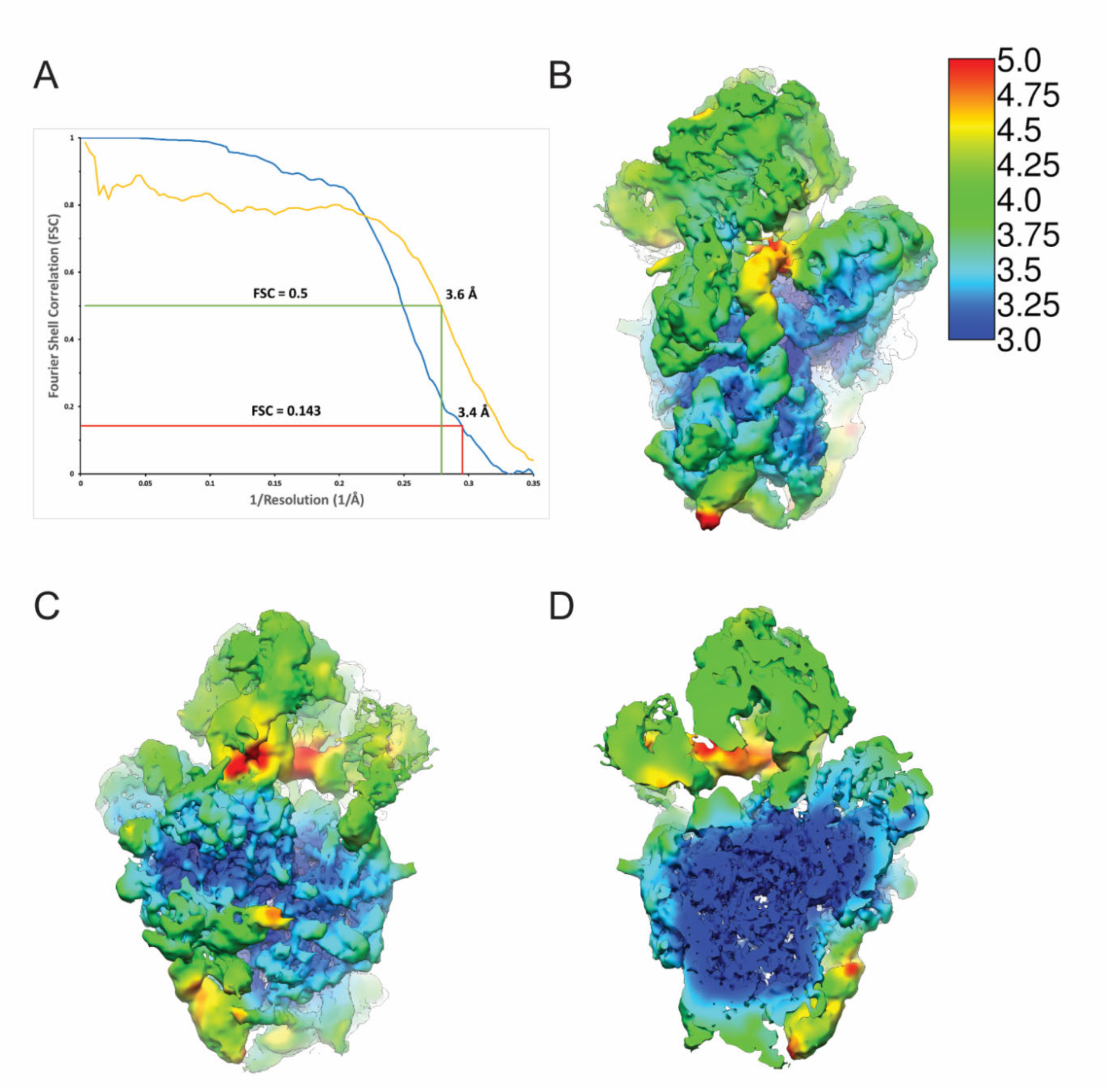
Local resolution heat map of the high-resolution pre-40S complex and comparison of the Fourier shell correlation curves calculated for the experimental 3D-EM reconstruction and the refined model of the pre-40S complex. **a.** Fourier Shell Correlation (FSC) curves for the high-resolution EM map (blue) and the one calculated from the model versus the map (yellow). At the FSC=0.143 criterion, the overall resolution of the high-resolution EM map was estimated to be 3.4 Å, which coincides well with the 3.6 Å resolution obtained for the model versus map FSC at a value of 0.5. **b-c.** Local resolution heat map two different orientations of the reconstruction of the full Pre-40S complex. The local resolution varies between 3 Å for the core of the ribosomal subunit and 4-5 Å for the peripheral parts of the rRNA expansion segments as well as the head of the Pre-40S and the assembly factors. Local resolutions were calculated using the “relion_postprocess” from Relion2.1 (Kimanius, Forsberg et al., 2016). **d.** Cut-through of the view in panel **b**.

**Appendix Figure S3:**
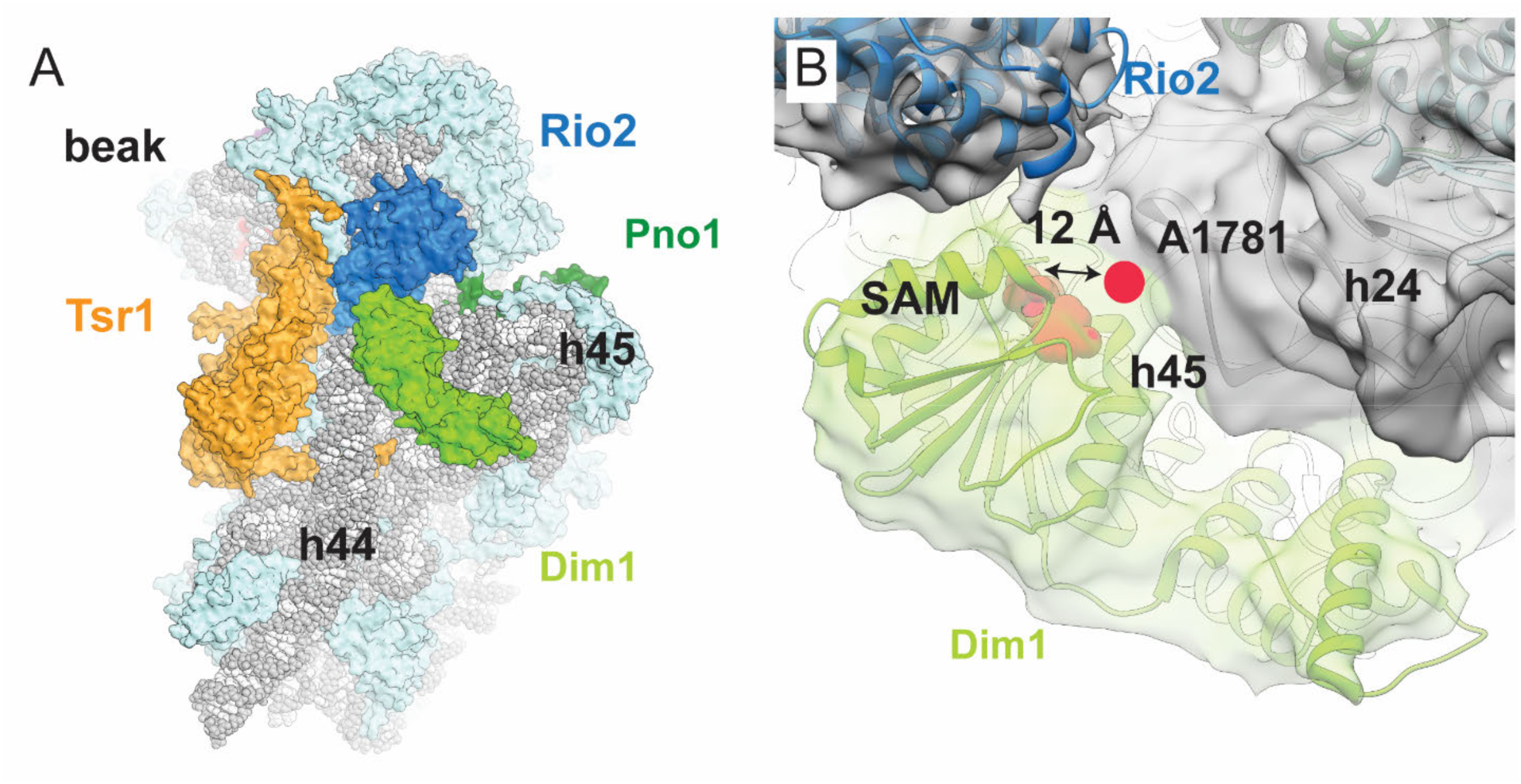
3D-reconstruction of the Dim1-containing subclass of cytoplasmic pre-40S particles. **a**. Overview over the Dim1-containing complex (r-proteins in light blue, rRNA in gray) from the intersubunit face. The rRNA methylase Dim1 (bright green) binds to the complex between the platform and the top of the displaced helix 44 in close vicinity of Rio2 (blue), Tsr1 (orange) and the decoding center. **b**. Dim1 (bright green) binds not only to helix 44, but also helices 45 and 24. Docking of the available crystal structure of human Dim1 in complex with S-adenosyl-methionine (SAM) (pink) (PDB : 1ZQ9) into the cryo-EM density shows that the previously described active site of Dim1 would be located too distant from its methylation target, A1781 (pink circle) to allow the methylation reaction to take place. Thus Dim1 has likely already methylated A1781 and A1782.

**Appendix Figure S4:**
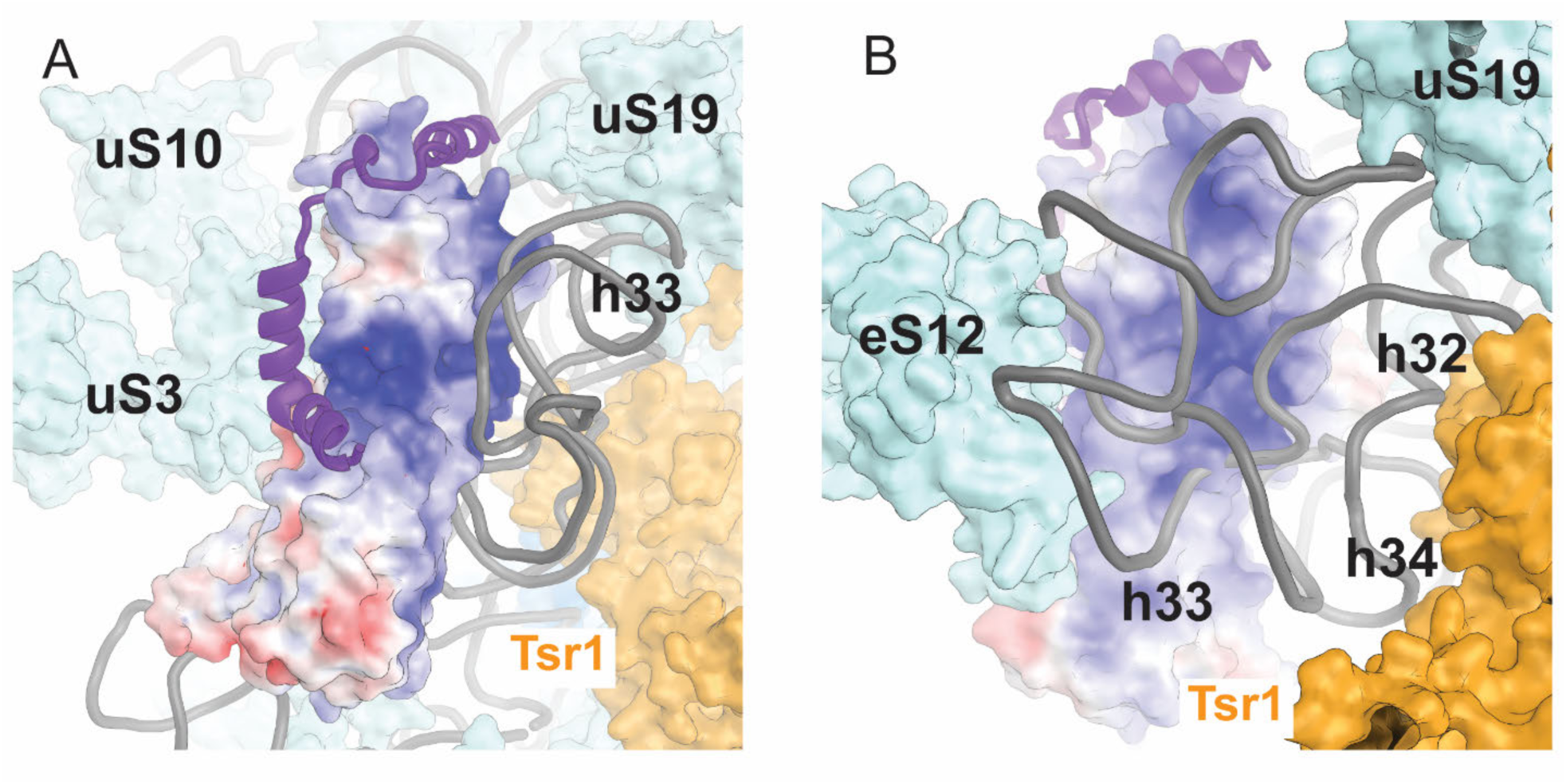
Surface representation of electrostatic surface charges of Enp1. **a-b.** Surface representation of Enp1 colored by electrostatic potential in context of the cytoplasmic pre-40S rRNA (gray), the r-proteins uS10, uS3, uS19 and eS12 (light blue), Tsr1 (orange), and Ltv1 (purple). The large postiviely charged surface (blue) is directly interacting with the backbone of the three-way junction of helices 32, 33 and 34. **a**. Front view from the tip of the beak. eS12 is hidden for clarity. **b**. Side view from the intersubunit space.

**Appendix Figure S5:**
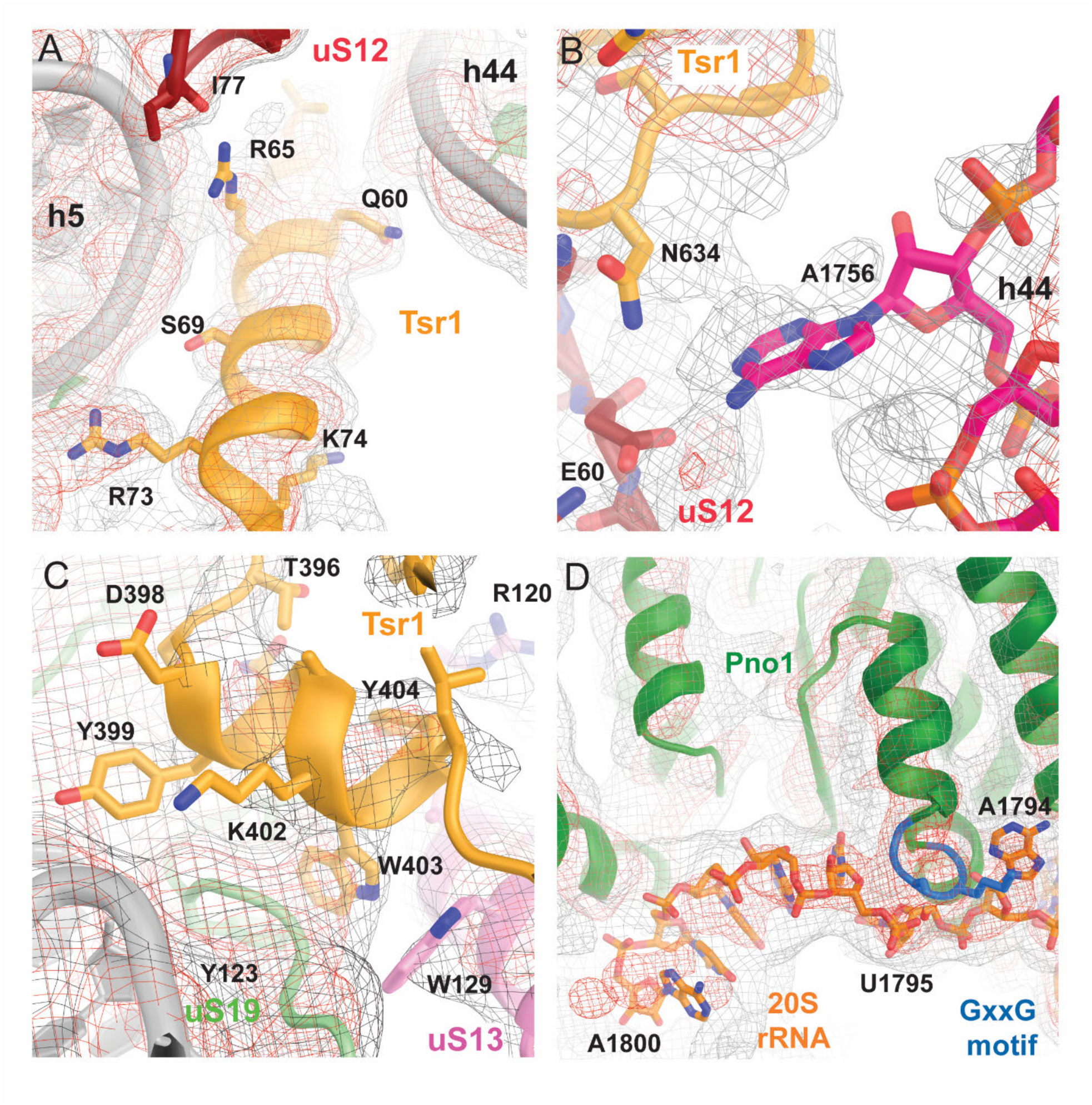
Details of the of functionally important assembly factors interactions in the cryo-EM map of the cytoplasmic pre-40S. **a-d.** The unfiltered cryo-EM map is shown at two different contour levels (gray and red mesh). **a**. N-terminal α-helix of Tsr1 (orange) interacting with helix 44. **b**. Visualization of the flipped out position of the decoding center base A1756 at the top of helix 44 (pink). **c**. Interactions of the conserved loop of Tsr1 with uS19 (green) and uS13 (pink). **d**. Interaction of the Pno1 (green) C-terminal KH domain (containing a GxxG RNA binding motif shown in blue) with the 3’terminal bases of the rRNA (orange).

**Appendix Figure S6:**
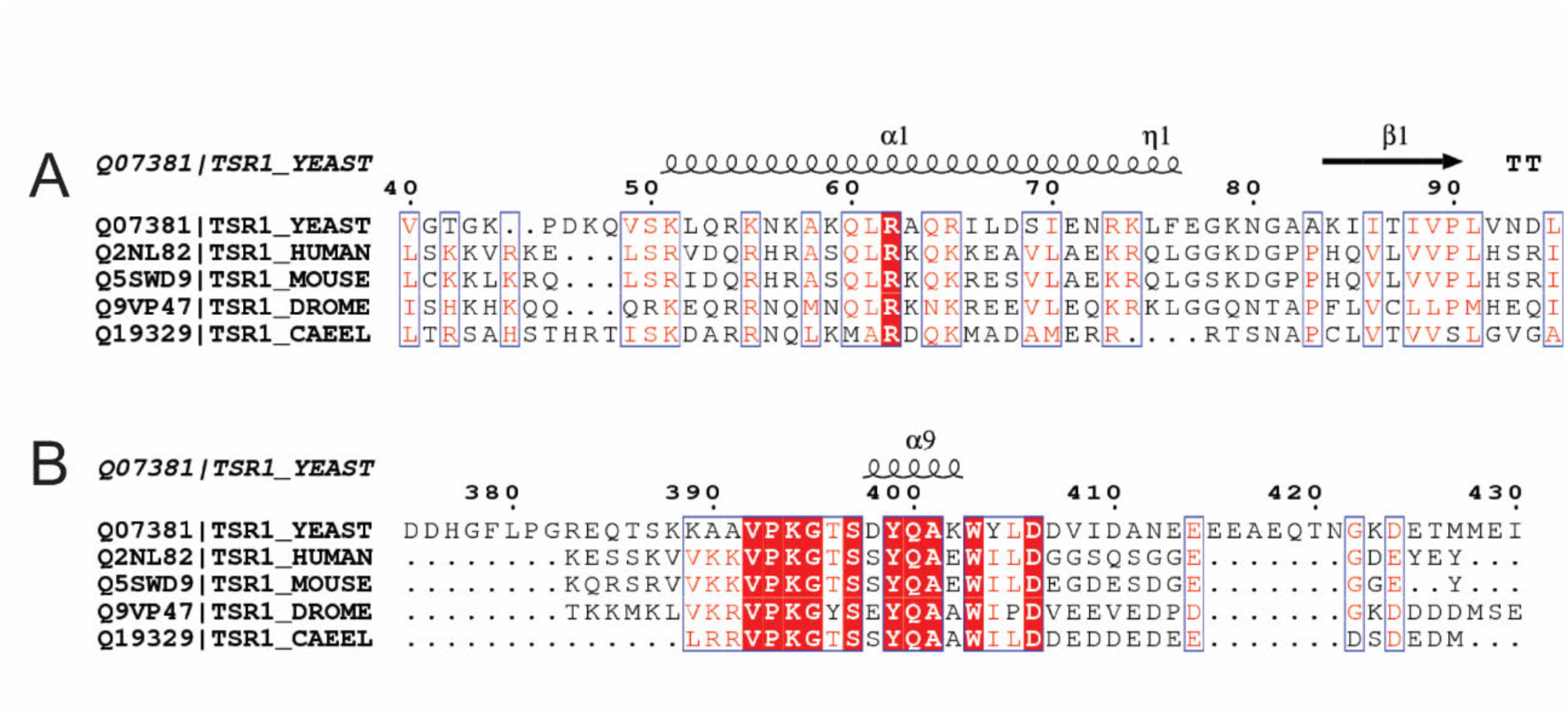
Multiple sequence alignments, conservation and secondary structure of parts of assembly factor Tsr1. **a**. Multiple sequence alignment of the N-terminus of Tsr1. Residues 51 to 75 (in *S. cerevisiae*) had not been resolved in the previously reported crystal structure of Tsr1 (McCaughan, Jayachandran et al., 2016), but appear as an α-helix in our cryo-EM map of the cytoplasmic pre-40S particle (as shown in Fig. 4b). The helix contains conserved positively charged residues (e.g. R62 and R73) that make contacts with the rRNA backbone of helices 5, 11 and 44. **b**. Multiple sequence alignment of the previously unresolved loop of Tsr1 (comprising residues 380 to 409 in *S. cerevisiae*). The cryo-EM map of the cytoplasmic pre-40S particle reveals that the conserved residues 392-408 specifically contact the displaced helix 31, as well as ribosomal proteins uS13 and uS19 and that residues 398-404 form a short α-helix (Fig. 4c).

**Appendix Figure S7:**
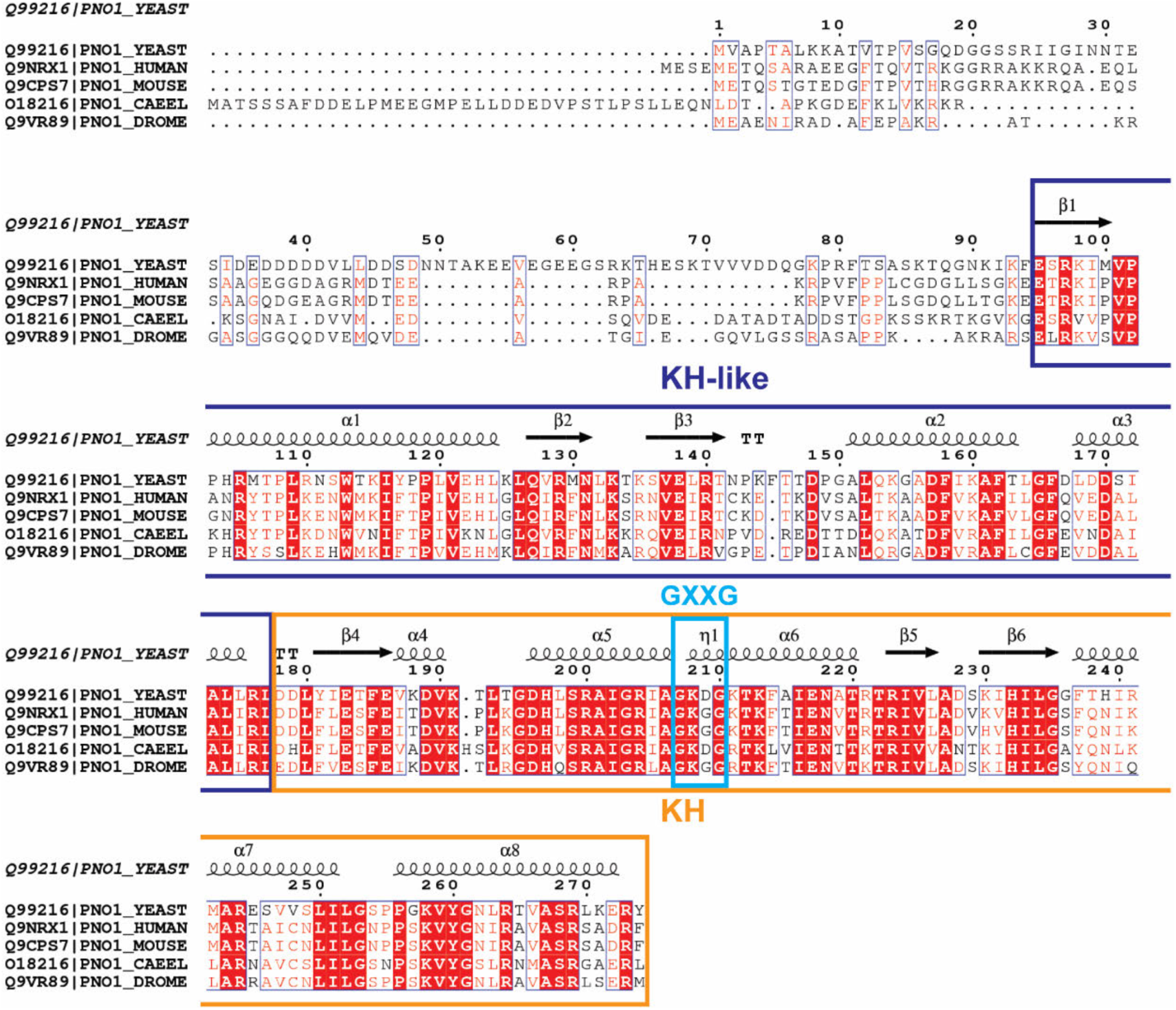
Multiple sequence alignment, conservation and secondary structure predictions of assembly factor Pno1. Multiple sequence alignment of Pno1 to highlight the high conservation of the KH-like (squared in blue) and KH domain (squared in orange). The poorly conserved N-terminal region was not resolve in our cryo-EM density. The conservation of Pno1 surface is shown in Figure 5c.

**Appendix Table S1:** Yeast strains used in this study.

| **Strain name** | **Genotype** | **Origin** |
| --- | --- | --- |
| BY4741 | *MATa ura3 his3 leu2 met15 TRP1* | Euroscarf |
| *P<sub>GAL1</sub>-NOB1* | *MATa ura3 his3 leu2 met15 TRP1 P<sub>GAL1</sub>-NOB1::natNT2* | This study |
| *P<sub>GAL1</sub>-HRR25* | *MATa ura3 his3 leu2 met15 TRP1 P<sub>GAL1</sub>-HRR25::natNT2* | This study |
| Enp1-GFP | *MATa ura3 leu2 TRP1 ENP1-GFP::HIS3MX* | Open biosystems |
| Tsr1-GFP | *MATa ura3 leu2 TRP1 TSR1-GFP::HIS3MX* | Open biosystems |

**Appendix Table S2:**
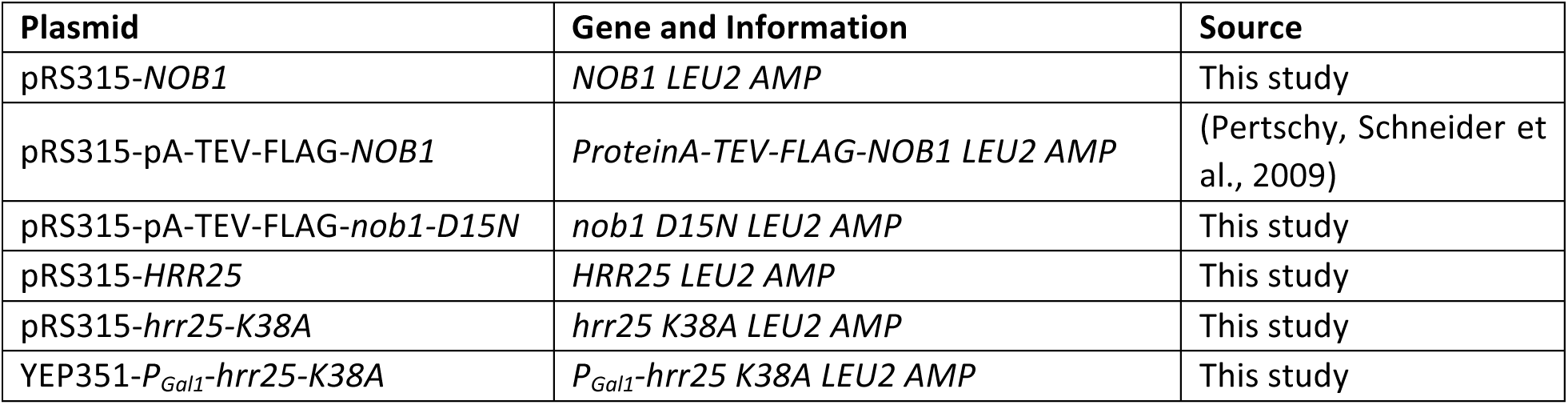
Plasmids used in this study.

| **Plasmid** | **Gene and Information** | **Source** |
| --- | --- | --- |
| pRS315-*NOB1* | *NOB1 LEU2 AMP* | This study |
| pRS315-pA-TEV-FLAG-*NOB1* | *ProteinA-TEV-FLAG-NOB1 LEU2 AMP* | (Pertschy, Schneider et al., 2009) |
| pRS315-pA-TEV-FLAG-*nob1-D15N* | *nob1 D15N LEU2 AMP* | This study |
| pRS315-*HRR25* | *HRR25 LEU2 AMP* | This study |
| pRS315-*hrr25-K38A* | *hrr25 K38A LEU2 AMP* | This study |
| YEP351-*P<sub>Gal1</sub>-hrr25-K38A* | *P<sub>Gal1</sub>-hrr25 K38A LEU2 AMP* | This study |

**Appendix Table S3:** Local resolution of the factors based on their atom position.

| **Factor** | **Chain ID** | **Resolution (relion_postprocess)** |
| --- | --- | --- |
| Pno1 | H | 3.6 |
| Enp1 | I | 4.1 |
| Ltv1 | J | 4.2 |
| Tsr1 | K | 3.7 |
| Rio2 | L | 4.0 |

**Appendix Table S4:** EM data collection and model statistics.

| **EM Data collection** |  |  |  |  |  |
| --- | --- | --- | --- | --- | --- |
| Number of datasets | 1 |  |  |  |  |
| Number of micrographs collected | 3,083 |  |  |  |  |
| Pixel size (Å) | 1.39 |  |  |  |  |
| Defocus range (µm) | 0.6 – 2.5 |  |  |  |  |
| Voltage (kV) | 300 |  |  |  |  |
| Electron dose (e<sup>-</sup>Å<sup>-2</sup>) | 40 |  |  |  |  |
| Name of 3D-reconstruction/model |  |  |  |  |  |
|  | Pre-40S complex | 80S-like particle | Dim1-containing particle | 3D reconstruction after focused classification on Enp1 | 3D-reconstructin after focused classification on Rio2 |
| EMDB entry of map | EMD-XXXX | EMD-XXXX | EMD-XXXX | EMD-XXXX | EMD-XXXX |
| PDB entry of the full model | PDB XXXX |  |  |  |  |
| Final number of particles | 165,168 | 145,081 | 40,044 | 39,367 | 61,562 |
| Resolution (Å) | 3.4 | 3.1 | 4.3 | 4.0 | 3.7 |
| Map sharpening B-factor (Å<sup>2</sup>) | -168.9 | -156.7 | -200.0 | -188.2 | -179.6 |
| **Refinement and model validation statistics<sup>(a)</sup>** |  |  |  |  |  |
| Resolution range ( Å) | 50-3.5 |  |  |  |  |
| Spacegroup | P1 |  |  |  |  |
| a=b=c (Å) | 278 |  |  |  |  |
| α=β=γ (°) | 90 |  |  |  |  |
| Number of reflections | 1'048'658 |  |  |  |  |
| Average B-factor (Å<sup>2</sup>) | 97.38 |  |  |  |  |
| Clashscore (all atoms) | 14.12 |  |  |  |  |
| R-factor (%) | 29.7 |  |  |  |  |
| wxc weighting value | 1.4 |  |  |  |  |
| Protein |  |  |  |  |  |
| Good rotamers (%) | 88.62 |  |  |  |  |
| Rmsd (bonds) | 0.007 |  |  |  |  |
| Rmsd (angles) | 1.023 |  |  |  |  |
| Ramachandran plot (%) |  |  |  |  |  |
| favored | 86.35 |  |  |  |  |
| allowed | 9.74 |  |  |  |  |
| outliers | 3.91 |  |  |  |  |
| RNA |  |  |  |  |  |
| Correct sugar puckers (%) | 96.6 |  |  |  |  |
| Good backbone conformation (%) | 65.1 |  |  |  |  |

<sup>(a)</sup>The model was validated using MolProbity implemented in PHENIX.REFINE (Adams, Afonine et al., 2010)

